# Epidemiology of *Legionella*: Genome-bAsed Typing (el_gato) - a new bioinformatic tool for identifying sequence-based types of *Legionella pneumophila* from whole genome sequencing data

**DOI:** 10.64898/2026.03.20.713011

**Authors:** Alan J. Collins, Dev Mashruwala, Vasanta Chivukula, Natalia A. Kozak-Muiznieks, Lavanya Rishishwar, Emily T. Norris, Melisa J. Willby, Jennafer A. P. Hamlin, Will A. Overholt

## Abstract

Sequence-based typing (SBT) via Sanger sequencing has been the standard for describing *Legionella pneumophila* relationships for two decades. SBT involves sequencing seven loci, identifying alleles using the United Kingdom Health Security Agency (UKHSA) database, and inferring the corresponding sequence type (ST). While similar SBT approaches for other organisms can be easily adapted to whole genome sequencing (WGS), *L. pneumophila* presents two known challenges for this adaptation: multiple copies of one locus (*mompS*) and extensive heterogeneity in a second locus (*neuA/neuAh*). Although several computational methods have been proposed to address these issues, a WGS-based replacement with equal resolution to traditional SBT has been elusive. To address this gap, we developed el_gato (**E**pidemiology of **L**egionella: **G**enome-b**A**sed **T**yping; https://github.com/CDCgov/el_gato), which offers several advantages over existing methods: (1) a novel approach for resolving multiple *mompS* alleles identified in the same isolate, (2) the ability to capture diverse *neuA/neuAh* alleles, (3) fast runtime with an average of 27.7 seconds per sample, (4) easy installation via Bioconda or Docker and (5) an updated database as of March 2025. el_gato works with either paired-end short reads or genome assemblies, performing more accurately with paired-end short reads at least 250 base pairs (bp) in length. We compared el_gato against two other *in silico* SBT tools (“mompS”, hereafter referred to as mompS tool and “legsta”) using a dataset of 441 isolates with sequence types (STs) previously determined by Sanger-based sequencing. el_gato correctly identified the ST for 98.9% of the test isolates, compared to 95.2% for the mompS tool and 42.2% for legsta, demonstrating a significant improvement compared to the mompS tool (adjusted p = 1.06e-3) and legsta (adjusted p = 4.24e-55) in ST identification. Furthermore, el_gato’s determination of ST was not significantly different from Sanger sequencing (adjusted p = 0.442). In summary, el_gato significantly improves *in silico* SBT and given its growing adoption, is poised to support the public health community.

## Introduction

*Legionella pneumophila* is a gram-negative bacterium that causes Legionnaires’ disease (LD), a severe pneumonia (McDade *et al*., 1977; Fraser *et al*., 1977; Brenner, 1979). It is commonly found in natural environments, such as lakes and soil, but can also thrive in human-made water systems, such as cooling towers and hot tubs (Fliermans *et al*., 1981; Ikedo and Yabuuchi, 1986; Euser *et al*., 2010; Buse *et al*., 2012; Dilger *et al*., 2018; Yao *et al*., 2024). When conditions are permissible for *Legionella* growth and there is a source of aerosolization, exposure may occur via inhalation resulting in LD (Davis *et al*., 1982; Bollin *et al*., 1985; Allegra *et al*., 2016). LD can occur as isolated cases, also known as sporadic cases, or as part of an outbreak comprised of cases related to a common source and identified within a specific timeframe. The U.S. Centers for Disease Control and Prevention (CDC) and the European Centre for Disease Prevention and Control (ECDC) have reported an increasing incidence of LD cases over the past two decades (Barskey *et al*., 2022; ECDC, 2022). Between 2015 and 2020 in the United States, *L. pneumophila* was the leading cause of drinking water–associated outbreaks, responsible for 786 illnesses, 544 hospitalizations, and 86 deaths and most often linked to premise plumbing in health care and lodging facilities (Kunz *et al*., 2024). Given these impacts, rapid identification and remediation of the source of infection is crucial.

Categorizing bacterial strains based on their genetic content provides multiple benefits. Of primary importance to public health, understanding the genetic relatedness of the pathogen supports effective epidemiological investigations in which outbreaks are detected, sources are identified, and control measures are implemented. Multi-locus sequence typing (MLST) is one of the most widely used molecular methods for bacterial characterization (Maiden *et al*., 1998; Urwin and Maiden, 2003; Maiden, 2006). For *L. pneumophila*, the MLST application is referred to as sequence-based typing (SBT) and analyzes seven genes (*flaA*, *pilE*, *asd*, *mip*, *mompS*, *proA*, and *neuA/neuAH*), to derive an allelic profile and ultimately a sequence type (ST). The United Kingdom Health Security Agency (UKHSA) *L. pneumophila* database is composed of nucleotide sequences associated with each previously identified allele for the seven genes in the typing scheme along with a list of STs derived from each allelic profile. As of March 2025, the number of defined alleles for each gene is as follows: *flaA*: n = 45, *pilE*: n = 66, *asd*: n = 90, *mip*: n = 96, *mompS*: n = 113, *proA*: n = 62, *neuA/neuAh*: n = 111 which make up 3241 allele combinations with an assigned ST. Allele profiles and their associated ST provide a convenient means to associate genetic relatedness between isolates of interest (Gaia *et al*., 2003, 2005; Ratzow *et al*., 2007; Lück *et al*., 2013).

While traditional SBT was developed using Sanger sequencing, the use of whole genome sequencing (WGS) has led to the development of bioinformatic tools that can derive SBT directly from paired-end short-read sequencing data. One major challenge for sequence typing with short-read sequencing data is identifying the correct allele when multiple copies of a target locus exist in the bacterial genome as, it can be difficult to assign reads correctly to each allele. In such cases, point mutations or insertions/deletions within each copy introduce variations that complicate typing accuracy (Li *et al*., 2008). The *mompS* gene is particularly problematic for *L. pneumophila* typing as multiple distinct copies are often present in the genome. Additionally, the *mompS* gene encodes a major outer membrane protein in *L. pneumophila* and is known to be under selective pressure for nucleotide variability (Gaia *et al*., 2003). In traditional SBT of *L. pneumophila* using Sanger sequencing, the typing copy of *mompS* used for ST determination, defined as *mompS2* but referred to here as *mompS*, is specifically amplified because the reverse primer (mompS1116R) binds downstream of the typing copy only (Gaia *et al*., 2005; Gordon *et al*., 2017, Figure 1). Since the ST depends on the allele of *mompS*, accurate identification of the typing copy is essential to ensure proper ST identification (Moran-Gilad *et al*., 2015; David *et al*., 2016; Raphael *et al*., 2016; Gordon *et al*., 2017).

**Figure 1:**
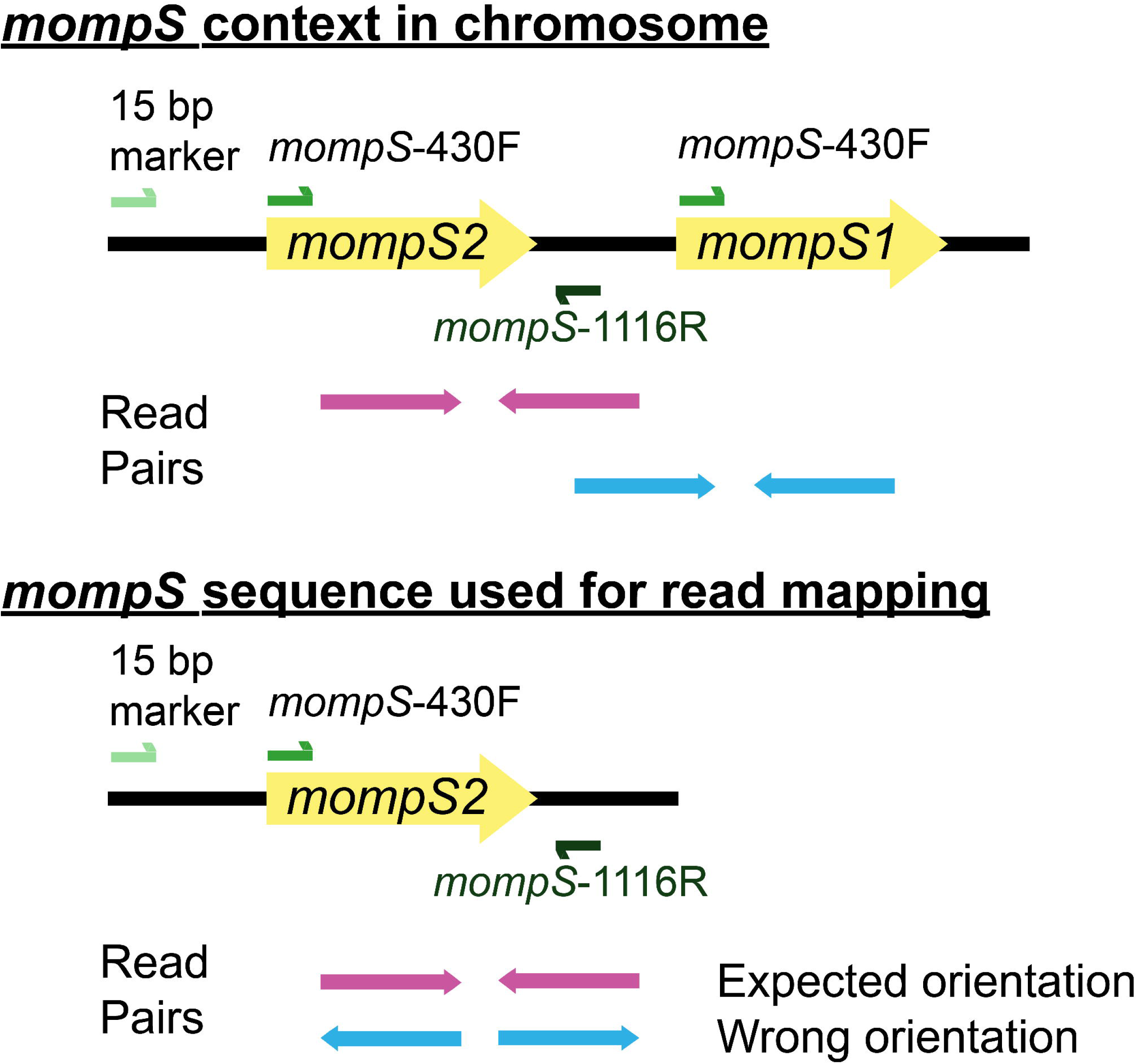
Resolving *mompS* alleles. a) Multiple copies of *mompS* may be found within *L. pneumophila*. Sanger SBT primers (mompS-430F and mompS-1116R) specifically target the typing copy based on the binding location of the reverse primer. In contrast when a WGS approach is used, read mapping programs cannot differentiate between the two copies using read mapping alone, potentially resulting in incorrect base assignment to *mompS*. b) el_gato uses the location and orientation of *momps*-1116R primer sequence, which is found downstream of only the typing *mompS* copy, to determine the correct typing copy. In addition, reads overlapping the 15 bp diagnostic marker described in Gordon et al (2017) are filtered. This figure is adapted from Gordon et al. (2017).

The *neuA* gene was incorporated to the SBT scheme a few years after the original scheme was released, as this gene was shown to increase the level of discrimination providing additional subdivision of common STs (Ratzow *et al*., 2007). *neuA* from the Paris strain (NC_006368.1) served as *neuA* allele 1; however, the *neuA* gene from the Paris strain presented additional challenges for *L. pneumophila* typing. After its inclusion in the SBT scheme, several studies reported isolates in which *neuA* could not be amplified using the standard primers. A common feature of these isolates was that they did not belong to serogroup 1; in other words, they were non-SG1 (Harrison *et al*., 2009; Amemura-Maekawa *et al*., 2010; Reimer *et al*., 2010; Vergnes *et al*., 2011). Farhat et al. (2011) showed that the *neuA* allele of the Dallas-1E strain (serogroup 5) of *L. pneumophila* shared at most 68% homology to the Paris strain *neuA* allele. In addition, some non-SG1 isolates carried a deletion that resulted in a 351 bp *neuA* fragment (allele 207) compared with the 354 bp amplicon in the Paris strain (allele 1) or the Dallas-1E strain (allele 201; Farhat *et al*., 2011). The role of *neuA* in lipopolysaccharide synthesis likely drives its high allelic diversity (Lüneberg *et al*., 1998; Zähringer *et al*., 1995) and this allelic diversity ultimately necessitated development of alternate SBT primers targeting *neuA* homologs (Mentasti *et al*., 2014). Alleles amplified exclusively by the alternate primers were classified as *neuAh* (*neuA* homolog). This variability also complicated WGS-based SBT, since mapping of sequencing reads and ultimately the identification of the *neuA/neuAh* allele relies on homology with a reference allele.

Several WGS-based tools, such as legsta (Seemann 2016) and the mompS tool (Gordon et al., 2017) have been developed to derive SBT from whole-genome data. *L. pneumophila* computational tools have incorporated steps to identify the typing copy of *mompS* gene specifically. For example, after mapping reads to the *mompS* reference sequence, the mompS tool retains paired-end reads that overlap regions specific to the typing copy at either end (15 bp marker or mompS1116R primer), ensuring that reads only associated with the typing copy of *mompS* are used for determination (Figure 1, Gordon *et al*., 2017). In contrast, another SBT computational tool, legsta (Seemann, 2016), is a wrapper around the *in silico* PCR (isPcr) tool (Kuhn *et al*., 2013). It takes as input a FASTA file and uses a primer sequence file, including *mompS* primers specifically designed to amplify the typing copy. Both legsta and the mompS tool include an allele and ST database specific to *L. pneumophila* and derived from the UKHSA database; however, they were last updated over five years ago and therefore do not reflect the most current *L. pneumophila* allele and ST database. In addition, while these tools implement targeted approaches to identify the typing copy of *mompS*, they do not incorporate a strategy to address the known heterogeneity of the *neuA*/*neuAh* locus, which could result in incomplete assignments for isolates with alternative *neuA/neuAh* loci.

This manuscript introduces el_gato - **E**pidemiology of **L**egionella: **G**enome-b**A**sed **T**yping, a bioinformatic tool designed to identify the ST of *L. pneumophila* from short read sequencing data. We evaluate a dataset of 441 isolates with previously generated Sanger-based SBT results and compare the ST calls between el_gato, legsta, and mompS tools. el_gato achieved higher accuracy than existing tools, runs quickly, and is easily installable as compared to the other tools. We show that el_gato outperforms other open-source tools for *L. pneumophila* ST determination and produces results comparable to Sanger-based methodology.

## Methods and Materials

### *L. pneumophila* data used during el_gato development

The dataset was composed of sequences generated from 441 isolates from the CDC *L. pneumophila* collection (Supplemental Table 1). These isolates were chosen as they have allele calls for all seven SBT loci previously determined by Sanger sequencing with an identified ST except for one isolate with a non-typed *neuA* (D4954). Sanger-based SBT was performed according to the European Society of Clinical Microbiology and Infectious Diseases Study Group for Legionella Infections (ESGLI) SBT protocol for typing of *L. pneumophila* with M13-tagged primers (Menstasti and Fry, 2012); however using mompS-1126R primer (Supplemental Table 2). In cases when *flaA* and *neuA* gene fragments failed to amplify with standard primers, the alternative flaA-L-N and flaA-R-N (Kozak-Muiznieks *et al*., 2018) and *neuAh* primer pairs were used (Mentasti *et al*., 2014)

Isolate collection years ranged from 1982 – 2017 and isolates came from 45 states, Canada (n = 1), France (n = 2), Denmark (n = 1), and two cruise ship investigations. The dataset consisted of 49 different STs, with multiple known problematic STs for which identification of the *mompS* allele and subsequent ST identification using previous *in silico* tools proved challenging due to the presence of duplicate non-identical *mompS* alleles (e.g., *mompS* 7 and 15; Krøvel *et al*., 2023). FASTQ data from this project is available on NCBI under Bioproject PRJNA936192.

### Sequence-based typing using el_gatov1.22.0 and Illumina reads

Nucleic acids were extracted from isolates using the Epicentre Masterpure DNA and RNA Purification kit (cat. No. MCD85201) per the manufacturer’s instructions. Illumina library preparation and sequencing used a protocol previously described (Mercante *et al*., 2018; Hamlin *et al*., 2024). Libraries were constructed using the Illumina Nextera XT Library Prep Kit (cat # FC-1311-1096) and the Nextera XT Index Kit v2 Set A (cat # VC-131-2001). Sequencing was performed using the MiSeq Reagent Kit v2 (500 cycles: cat # MS-102-2003) on the Illumina MiSeq.

For five of seven target loci (*flaA*, *pilE*, *asd*, *mip*, and *proA*) the process of allelic identification by el_gato is straightforward. Paired end reads are mapped to the *L. pneumophila* Paris strain (NC_006368.1) using minimap2 v2.28 (Li, 2018) to identify locus-specific reads. These reads are then used to determine the closest allele match within the included allele database, using BLASTn v2.16.0+ (Camacho *et al*., 2009) and applying specified thresholds (detailed below). el_gato default parameters are as follows: base calls with Phred quality scores below 20 are ignored; after filtering low-quality base calls in reads, any positions with depth of less than 10 (‘--depth’) are excluded; a default kmer size of 21 is used in minimap2 v2.28 (‘--kmer-size’); the required percentage of reference bases identified through mapping is set to 100% for six of the seven genes, for *neuA/neuAh* the value is 99% due to the 3 bp deletion present in certain alleles. All default parameters except the Phred quality score are adjustable using their associated flags. *mompS* and *neuA/neuAh* require additional processing steps (detailed below) prior to BLASTn identification of the closest allele match.

### Processing neuA/neuAh using el_gatov1.22.0 for sequence-based typing

The three *neuA/neuAh* alleles from three previously identified allele groups were used as references for mapping: neuA_1 allele from the Paris strain (NC_006368.1), the neuA_201 allele from the Dallas-1E strain (NZ_CP017458.1) and, neuA_207 (NZ_CP017601.1), which contains a 3 bp deletion relative to the first two alleles. During el_gato development, sequence divergence between *neuA* and *neuAh* references prevented some isolates from aligning to these three reference alleles. To mitigate this limitation and anticipate future divergent alleles, the phylogeny of neuA/neuAh was assessed using a minimum spanning tree based on pairwise sequence mismatches (Zhou *et al*., 2018; Figure 2).

**Figure 2:**
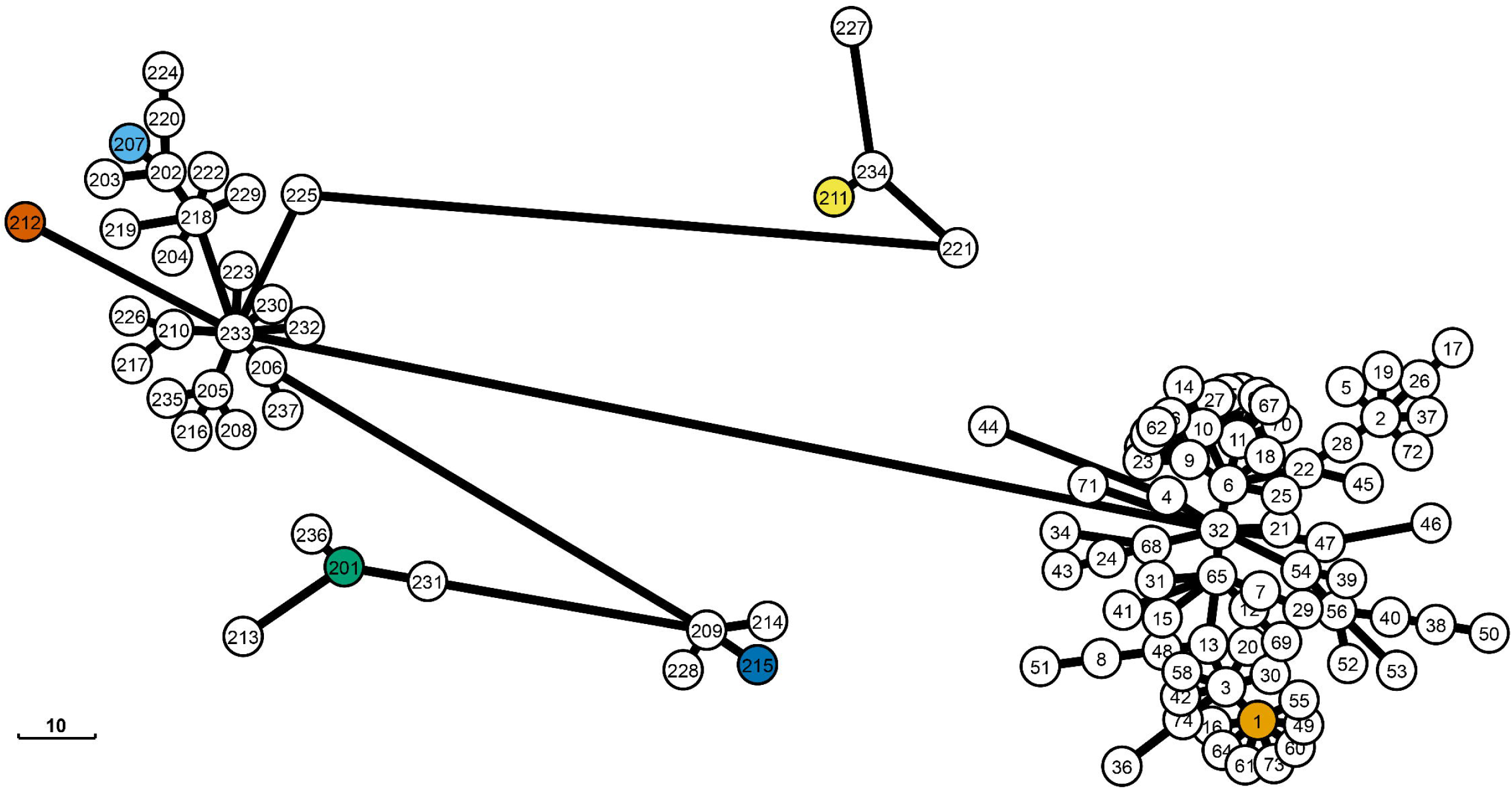
*neuA/neuAh* Sequence Diversity. The figure shows a minimum spanning tree based on nucleotide similarity of all *neuA/neuAh* alleles found within the el_gato database. Lines and the scale bar indicate the number of single nucleotide polymorphisms (SNPs). Three allele groups that encompass the known genetic heterogeneity in *neuA/neuAh* were previously identified (allele group Paris, neuA/neuAh_1; allele group Dallas-1E, neuA/neuAh_201; and neuA/neuAh_207). We identified three additional allele groups (neuA/neuAh_211, neuA/neuAh_212, and neuA/neuAh_215). References from all allele groups should be included for mapping short read sequencing data to best capture the breadth of *neuA/neuAh* diversity. The colored circles in each allele group indicate the reference alleles used for mapping.

This analysis identified three additional allele groups, and reference alleles corresponding to these allele groups were added: neuA_211 (KR350686.1), neuA_212 (GCA_015966455.1), and neuA_215 (GCA_030724125.1) which capture a broader range of *neuA/neuAh* genetic diversity. In el_gato, sequences are mapped to all six references to improve detection of more divergent *neuA/neuAh* alleles. The reference with the highest mapping depth is selected, provided that its depth is at least three times greater than that of any other reference allele, as determined by samtools (Danecek *et al*., 2021). If no query sequence meets the required coverage percentage against at least one *neuA/neuAh* reference, or if the top reference fails to meet the threefold depth criterion, no allele is assigned to the *neuA/neuAh* locus.

### Processing mompS using el_gato v1.22.0 for sequence-based typing

*mompS* is sometimes present in multiple distinct copies in *L. pneumophila* (Gaia *et al*., 2005; Gordon *et al*., 2017). Traditional Sanger-based methods overcome this by using a reverse primer specific for the typing copy of *mompS* during amplification (Gaia *et al*., 2005; Gordon *et al*., 2017). However, gene duplication poses a significant challenge for a short-read sequencing approach (Krøvel *et al*., 2023) where reads may not be accurately mapped. el_gato addresses this issue by leveraging the genomic proximity of the two *mompS* copies, which are ~2100 bp apart. The typing copy lies immediately upstream of the non-typing copy and is flanked by the primer-binding regions used in conventional Sanger-based SBT. While the forward primer (mompS-430F) binding site is found upstream of both copies, the reverse primer (mompS-1116R) binding site is present only downstream of the typing copy (Figure 1). By leveraging this orientation, el_gato distinguishes between the two copies ensuring that the typing *mompS* copy is identified. Specifically, the process for resolving the sequence of the two *mompS* copies and identifying the correct allele is as follows:

1. **Mapping Reads:** Reads are mapped to the reference *mompS* sequence (*L. pneumophila* Paris strain, NCBI NC_006368.1) and those aligning to either the typing or non-typing copy of *mompS* are retained.
2. **Identifying Biallelic sites:** el_gato examines the nucleotide sequence for each position within reads mapped to *mompS*. If more than 30% of the reads show base heterogeneity at a position, it is considered biallelic. When no biallelic positions are detected, el_gato extracts the *mompS* sequence and uses BLASTn to determine the allele number. In contrast, the presence of biallelic positions indicates multiple distinct *mompS* copies. In this case, el_gato identifies individual read pairs that map to biallelic positions and uses them to reconstruct the sequence of each possible allele.
3. **Identifying the Correct Allele:** el_gato identifies the typing *mompS* by analyzing read orientation relative to two diagnostic sequences flanking the locus. The first is a 15 bp marker upstream of the typing copy (5’-GTTATCAATAAAATG-3’), previously described by Gordon et al. (2017), which is absent in the non-typing copy. Read pairs containing this marker can therefore be assigned to the typing copy. The second marker, the mompS-1116R primer sequence, lies between the two *mompS* loci and is downstream of the typing copy but upstream from the non-typing copy. The orientation of reads containing the mompS-1116R primer sequence, relative to the typing copy, is used to infer which copy is the typing locus (Figure 1). For read pairs where one or both reads map to at least one biallelic site, el_gato applies the following rules to assign reads to the typing locus:

I. Only one allele has reads with mompS-1116R sequence in the expected orientation (3’-5’).
II. One allele has >3x as many reads with mompS-1116R primer sequence in the expected orientation (3’-5’) compared to the other allele.
III. One allele has no reads with mompS-1116R primer sequence in either orientation, while the other allele has reads with mompS-1116R sequence in the incorrect orientation (5’-3’). In that case, the allele without the mompS-1116R sequence is considered the typing locus.
IV. Reads from one allele contain the upstream 15 bp diagnostic marker (at the start of the 987 bp *mompS* reference sequence) in the expected 5’ to 3’ orientation, while the other allele does not. These reads act as secondary evidence that the allele represents the typing copy.
V. If no reads contain either the mompS-1116R primer sequence with the correct orientation or upstream 15 bp diagnostic marker, el_gato cannot identify the typing locus.

### Processing isolates for analysis with alternative SBT tools

We performed ST determination with two open-source tools for *L. pneumophila* typing: mompS tool (https://github.com/bioinfo-core-BGU/mompS) and legsta (https://github.com/tseemann/legsta) in addition to el_gato. As both tools require assemblies, we processed the 441 isolates using standard bioinformatic methods for *de novo* genome assembly. Specifically, we trimmed reads using fastp v0.23.4 (Chen *et al*., 2018) with default settings. We performed *de novo* assembly with spades v3.15.3 (Prjibelski *et al*., 2020) with default settings and the ‘--careful’ flag generating assemblies in FASTA format. Once completed, we filtered assemblies to remove contigs that were <1000 bp and summarized genome metrics using Quast v5.2.0 (Gurevich *et al*., 2013). Additionally, for both raw and trimmed read data used with el_gato, we used FastQC to assess read quality (Andrews, 2010). As the mompS tool requires both reads and assemblies, all input files were provided and run with default settings. In contrast, legsta was run with only assemblies and its default settings. el_gato v1.22.0 was run using reads as input and with default settings which include a minimum mapping depth of 10 reads per position and a kmer size of 21 when using minimap2. Finally, all tools were run single-threaded during all comparisons.

### Sequence-based typing using el_gatov1.22.0 and genome assemblies

In addition to read-based analysis, el_gato can also process assembled genomes directly; however, caution is warranted, as crucial information such as mapping quality and read orientation, is lost during the assembly process. Further details on how el_gato processes assemblies and associated results are provided in the supplemental materials.

## Results

Following sequencing, we first examined key quality metrics from the raw and trimmed data, along with genome assembly statistics. Raw sequence data collected for each isolate had an average of 1185148 reads (min: 105454 – max: 4876842), with an average read length of 244 bases (min: 149 – max: 251), and average sequencing coverage of 84X (min: 8 – max: 285) as estimated by number of reads times read length divided by the *L. pneumophila* genome size (~3.4 Mb; Supplemental Table 3). Trimmed sequencing data had an average of 1179639 reads (min: 105402 – max: 4862494), with an average read length of 243 bases (min: 149 – max: 250), and an average coverage of 83X (min: 8 – max: 285). Genome assembly metrics for isolates included an average number of contigs ≥ to 1000 bp of 33 (min: 9 – max: 95), an average total length of sequence 3, 4705, 732 bp (min: 17591 – max: 3831085), and an average N50 as 272, 332 bp (min: 1228 – max: 849757; Supplemental Table 4).

The accuracy for each platform (el_gato, legsta, and mompS tool) was evaluated for 441 isolates, using Sanger sequencing derived ST calls as the comparator. We also assessed gene-level accuracy within the 7-gene scheme. Because the mompS tool and legsta require assemblies, which were generated after quality control with fastp (Chen *et al*., 2018), we compared el_gato accuracy on read data with and without fastp processing (Supplemental Table 5). To determine the accuracy, we used the following formula to calculate percent accuracy:

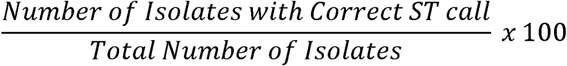

### el_gato v1.22.0 Accuracy

el_gato demonstrated no differences when using raw sequencing data or data processed with fastp (Supplemental Table 5). el_gato correctly predicted 98.9% ST calls (n = 436/441). Correctly identified alleles per locus, as compared to the Sanger results were as follows: *flaA* 100% (441/441), *pilE* 100% (441/441), *asd* 99.8% (440/441), *mip* 100% (441/441), *mompS* 99.5% (439/441), *proA* 99.8% (440/441), and *neuA/neuAh* 99.8% (440/441; Supplemental Table 5). There were no incorrect calls made for el_gato, only instances where no ST calls were made due to an incomplete allele profile (discussed below).

### legsta v0.5.1Accuracy

legsta correctly predicted 42.2% (186/441) of ST calls. Correctly identified alleles per locus, as compared to the Sanger results were as follows: *flaA* 89.6% (395/441), *pilE* 96.6% (426/441), *asd* 98.6% (435/441), *mip* 98.4% (434/441), *mompS* 49.9% (220/441), *proA* 98.6% (435/441), and *neuA/neuAh* 98.6% (435/441; Supplemental Table 5). It incorrectly typed 57 isolates relative to Sanger sequencing. Of these, 54 isolates had complete allele profiles but could not be assigned an ST because the corresponding allele combinations are absent from both the legsta database and the 2025 database. For 51 isolates, the results were associated with *mompS* alleles 18 and 63. In addition, *mompS* was incorrectly called as *mompS* 7 for two isolates (D7508 and F1010), when Sanger identified it as *mompS* 15, and for isolate D7051, *mompS* 78 was determined instead of *mompS* 19. Finally, incorrect full allele profiles were determined and, as a consequence, incorrect STs were assigned to three isolates (D3851, D5746, and D7025).

### mompS Tool Accuracy

The mompS tool correctly predicted 95.2% ST calls (420/441). Correctly identified alleles per locus, as compared to the Sanger results were as follows: *flaA* 95.2% (420/441), *pilE* 95.2% (420/441), *asd* 95.2% (420/441), *mip* 95% (419/441), *mompS* 95.2% (420/441), *proA* with 95.2% (420/441), and *neuA/neuAh* 95.2% (420/441; Supplemental Table 5). There were no incorrect calls made by the mompS tool, only instances where no ST calls were made due to an incomplete allele profile (discussed below).

### Statistical analysis of all methods compared to Sanger’s results

To evaluate the performance of different ST identification methods, we first compared three approaches, el_gato (using raw FASTQ data), legsta, and mompS, to Sanger-identified STs for all isolates. We applied Cochran’s Q test using the rstatix v0.7.2 package in R v4.4.0 (R Core Team, 2021), where the binary outcome is either correctly called ST or incorrectly called ST. Cochrans Q test revealed significant differences among these methods (Q = 703, df = 3, *p* = 4.80 × 10□^1^□^2^) and indicates that at least one of the methods differs in ST assignment accuracy. We also compared el_gato outputs generated from raw or trimmed reads and assembled genomes as well as results from legsta and the mompS tool. This comparison confirmed significant differences across all six methods (Q = 1004, df = 5, p = 1.01 × 10□^21^□).

To determine which method(s) differed statistically from Sanger, we performed pairwise McNemar’s tests using the rstatix v0.7.2 package in R v4.4.0. First, we demonstrated that read trimming had no effect on el_gato results: analyses using either raw or trimmed FASTQ data produced ST assignments that were statistically indistinguishable from those obtained by Sanger sequencing (adjusted p = 1, Supplemental Table 6). As there was no significant difference between the results from raw and trimmed FASTQ data, trimming had minimal impact on ST call accuracy (Supplemental Table 6). el_gato with either raw or trimmed inputs was significantly different from the other WGS tools (legsta: adjusted p = 9.90e-55; mompS tool: adjusted p = 2.48e-3), including el_gato using assemblies (adjusted p = 3.78e-38). Typing results generated from el_gato using assemblies were also significantly different from both legsta and the mompS tool methods (adjusted p = 3.72e-14 and 3.01e-32 respectively; Supplemental Table 6). While el_gato was not significantly different from Sanger sequencing (adjusted p = 1.00), the other computational tools were significantly different from Sanger sequencing (legsta: adjusted p = 8.05e-56; mompS tool: adjusted p = 1.78e-4).

### Tool comparison: ease of installation

The installation process varies between the three tools, presenting challenges depending on the user’s familiarity with the command line. For example, el_gato can be installed by either cloning from GitHub or installing via pip, though these require resolving several dependencies. To simplify this, el_gato and its dependencies are also distributed through Bioconda or Docker. legsta offers similar ease of installation with options for downloading from conda, Homebrew, or GitHub. In contrast, installing the mompS tool is more complex as once it’s downloaded from GitHub, users must manually install five dependencies and update the configuration file with the correct paths for each dependency.

### Tool comparison: database differences

Having an updated database for determining each allele profile and the associated ST is essential. The UKHSA database is continually updated as new alleles and STs are identified. The database distributed with el_gato, kindly provided by UKHSA in March 2025, will be updated at least annually. In contrast, the legsta database reflects the UKHSA database as it was five years ago. The mompS tool reflects the database from nine years ago. The total number of alleles in each tool’s database is as follows: el_gato – 598, legsta – 524, and the mompS tool – 460. Similarly, the number of ST profiles is as follows: el_gato – 3, 342, legsta – 2, 793, and the mompS tool – 1, 677.

Using legsta, 51 isolates could not be assigned an ST, as their allele combinations were not represented by any assigned ST. Furthermore, comparison with the el_gato database showed that these profiles are still not associated with a defined ST. Upon review, every one of these 51 isolates had the same incorrect call of mompS_63 instead of the correct mompS_18 allele. Although allele profiles can still be generated as long as all alleles have been previously reported, the reduced number of ST profiles limits the ability to assign STs, assess genetic relatedness, and communicate results effectively without the additional work of submitting the profile and Sanger sequencing output to the UKHSA database for allele assignment.

### Tool comparison: run time

The three tools demonstrated distinct runtime profiles. el_gato, using raw FASTQ data, averaged 27.43 seconds per sample (range: 6.00–89 seconds), el_gato using trimmed FASTQ data averaged 26.88 seconds per sample (range: 5.98–89.55 seconds), legsta averaged 3.11 seconds per assembly (range: 0.135–5.52 seconds), and the mompS tool averaged 366.92 seconds (~6 minutes) per sample (range: 35.6–1239.43 seconds; supplemental Table 1).

### Tool comparison: generated output

el_gato analysis results in the creation of seven output files in the output directory. Those files include: identified_alleles.fna, intermediate_outputs.txt, possible_mlsts.txt, reads_vs_all_ref_filt_sorted.bam, reads_vs_all_ref_filt_sorted.bam.bai, and run.log, which are detailed in the supplemental materials and on GitHub. Meanwhile, the ST and its associated allele numbers are reported as standard out when running the tool. If an ST cannot be found, the ST is recorded as an alternative symbol dependent on the results. For example, if data does not pass the default depth requirement of 10 for a particular locus, then a ‘-’ is reported for that allele, and subsequently ‘MD-’ (missing data) is reported for that ST. Further descriptions for all symbols are found on GitHub (https://github.com/CDCgov/el_gato) and in the supplemental materials. The generated bam file can be viewed using a tool such as integrated genome viewer (IGV; Thorvaldsdóttir *et al*., 2013) to review the mapping output in cases where el_gato is unable to generate a call.

Output from legsta does not indicate reasons why an allele could not be determined. The tool only reports the ST with allele numbers to standard output and does not produce any intermediate files. If an ST cannot be determined, the output shows either a ‘-’ to indicate “no in silico product detected” for an allele or a ‘?’ for a novel allele, which results in a ‘-’ for the ST.

The mompS tool generates 21 output files, mostly from mapping pipeline steps, along with outputs detailing how the *mompS* allele was determined. These files include SAM/BAM files (including deduplicated, filtered, and indexed versions), a consensus FASTA sequence with logs and BLAST indexes, and outputs from variant calling and typing analyses, including MLST, *mompS* typing, and BLAST searches. However, when an isolate fails to produce a full allele profile, there is no explanation provided as to why this occurred.

## Discussion

The application of SBT in *L. pneumophila* has been widely explored in the context of outbreak investigations and disease surveillance. Accurate ST calls are crucial for understanding the genetic relatedness during outbreaks (Gaia *et al*., 2003). As laboratories transitioned from Sanger sequencing to WGS-based SBT, certain limitations have slowed the full adoption for *L. pneumophila*. Here, we demonstrate the successful application of el_gato, a new short-read sequencing SBT algorithm for *L. pneumophila.* Our pipeline builds upon prior studies (Gaia *et al*., 2005; Gordon *et al*., 2017; Krøvel *et al*., 2023), improving accuracy for *mompS* and *neuA/neuAh* typing, generating user-friendly and customizable report documents, and offering rapid run time with simple installation.

Certain isolates remained non-typeable across all tools, underscoring differences among tools and the critical role of careful sequencing practices (Table 1). We investigated isolates for which el_gato, legsta, and the mompS tool all failed to generate an ST call. These included samples with short reads, small insert sizes, and low genome coverage. Short reads and small insert sizes posed challenges for allele determination in several isolates. For example, in D7119 and F4185, the base at position 509 of the typing *mompS* locus is biallelic, representing either a novel allele type (NAT) or *mompS_28* allele. Because the reverse primer spans positions 940–960, inserts or fragments must be >430 bp to reach the start of the mompS-1116R primer sequence or and the biallelic site. el_gato was unable to identify reads covering this region or spanning the bi-allelic site and the 15 bp diagnostic marker upstream of *mompS*, resulting in an ST call flagged as having multiple alleles (MA?). This was likely due to shorter average read lengths: 149 bp for D7119 and 150 bp for F4185. Additionally, small insert sizes (224 bp for D7119 and 222 bp for F4185) restricted paired-end read coverage across biallelic sites. In comparison, neither legsta nor the mompS tool provided possible allele information nor could generate an ST call for these isolates. Sanger sequencing confirmed the correct allele as *mompS_*28. As such, we are tracking alleles that are problematic for el_gato: https://github.com/CDCgov/el_gato/blob/main/docs/problematic_alleles.md.

**Table 1.** Additional information for samples that did not produce a ST call for el_gato.

| Sample ID | el_gatov1.22.0 missing allele calls | mompS tool missing allele calls | legsta missing allele calls | Genome coverage | Average Read Length (bp) |
| --- | --- | --- | --- | --- | --- |
| D7119 | mompS | mompS | flaA, pilE, & mompS | 59X | 149 |
| D7856 | neuA | all loci | mompS | 17X | 241 |
| F3359-001 | proA | all loci | all loci | 8X | 246 |
| F3359-003 | asd | all loci | all loci except neuA | 30X | 247 |
| F4185 | mompS | all loci | flaA, pilE, & mompS | 215X | 150 |

Low genome coverage also affected allele calling. For F3359-001 and D7856, coverage fell below el_gato’s required 10× depth at certain loci. Even though read length and coverage were marginally sufficient, F3359-003 failed to generate a call for *asd* via el_gato because most reads covering the 473 bp target region spanned only ~250 bp, preventing allele determination. These challenges are not unique to el_gato. Krøvel et al. (2023) reported that short reads, sequencing kit choice, and assembly method can affect *L. pneumophila* typing using legsta or BLASTn, with reads shorter than 300 bp sometimes causing misalignment or misassembly, particularly in the *mompS* gene. When an allele is not identified using *in silico* SBT, traditional SBT can be performed to resolve that allele or isolates can be re-sequenced and re-analyzed with el_gato.

While the mompS tool has been widely used for *L. pneumophila* SBT, it has known limitations (Krøvel *et al*., 2023). In our dataset, for example, it could not correctly type any ST1335 (n = 3) or ST1395 (n = 5) isolates, both belonging to *L. pneumophila* subspecies *pascullei* (Brenner, 1979; Kozak-Muiznieks *et al*., 2016, 2018). It also failed for three of 24 ST213 isolates and one of 48 ST222 isolates, due to the biallelic nature of the *mompS* gene in isolates with these STs. Additionally, ST determination with mompS tool failed when the maximum contig length in assemblies was below ~3, 200 bp (n = 6).

In el_gato, allele sequences are reconstructed using fixed reference coordinates. Insertions relative to the reference are ignored and deletions result in reduced coverage or unresolved positions. As a result, isolates containing indels may produce unexpected results. The legsta and the mompS tools also do not handle indels. Although indels are supported in *neuA*/*neuAh* we did not observe any indels in the other loci within this diverse dataset. Even so, using a large and diverse isolate collection, el_gato demonstrates robust ST calling, even for problematic *mompS* alleles. Users can easily generate informative reports, and with an updated database, el_gato is well positioned as the preferred tool for typing *L. pneumophila* from paired end short reads.

## Disclaimer

This research received no specific grant from any funding agency in the public, commercial, or not-for-profit sectors. The use of trade names is for identification only. It does not constitute endorsement by the U.S. Department of Health and Human Services, the U.S. Public Health Service, or the Centers for Disease Control and Prevention. The findings and conclusions in this report are those of the authors and do not necessarily represent the official position of the Centers for Disease Control and Prevention.

## Supporting information

SupplementalFigure1-elgatoBatchReport-Reads

SupplementalFigure2-elgatoBatchReport-Assemblies

SupplementalTable1-egMetaData

SupplementalTable2-SBT_primers

SupplementalTable3-RawAndTrimmedQualityMetrics

SupplementalTable4-QuastAssemblyMetricsAll

SupplementalTable5-egToolComparisonToSangerResults

SupplementalTable6McNemarOutput

## Acknowledgements

We gratefully acknowledge the contributions of all past members of the Legionella research team, whose efforts laid the foundation for this work including J. Caravas, A. Gaines, M. Ishaq, J. Mercante, S. Morrison, B. Raphael, and J. Winchell. We also kindly acknowledge the support of the UK Health Security Agency including B. Afshar, R. Tewolde, and S. Platt.

## Data Availability

All FASTQ data from this project is available on NCBI at Bioproject PRJNA936192.

## Supplemental Table Headers

**Supplemental Table 1:** Meta data for the isolates that were used during the development of el_gato. Column headers are as follows:

sampleName – identifier of each isolate

year – year when the isolate was collected or unknown

source – source of the isolate: clinical sporadic (CS), clinical outbreak (CO), environmental isolate (EN), or environmental outbreak (EO)

state – geographic location where the isolate was collected, or unknown (N/A) or cruise ship (INT).

timeToCompleteInSeconds-eg1.22.0-Raw – amount of time el_gato took to complete per isolate

timeToCompleteInSeconds-eg1.22.0-Trimmed – amount of time el_gato took to complete per isolate when sequences had been previously processed via fastp for quality control

timeToCompleteInSeconds-legsta – amount of time legsta took to complete per isolate

timeToCompleteInSeconds-mopmS – amount of time the mompS tool took to complete per isolate

timeToCompleteInSeconds-elgato1.22.0Assemblies - amount of time el_gato took to complete per isolate assembly

**Supplemental Table 2:** PCR amplification and sequencing primers used during Sanger sequence-based typing for the isolates examined in this study. The bold text indicates M-13 tags added to amplification primers.

**Supplemental Table 3:** Fastqc output on the read quality for all isolates used in this study. Column headers are as follows:

sampleName – identifier of each isolate

numberOfRawReads – count of reads before fastp trimming for each isolate

avgRawReadLength-bp – average read length before fastp trimming for each isolate

rawCoverage – genome coverage calculation as number of reads times read length divided by *L. pneumophila* genome size of 3.4 Mb without performing any quality control steps via fastp

numberOfTrimmedReads – count of reads after fastp trimming for each isolate

avgTrimmedReadLength-bp – average read length after fastp trimming for each isolate

trimmedCoverage – genome coverage calculation as number of reads times read length divided by *L. pneumophila* genome size of 3.4 Mb after performing quality control steps via fastp

**Supplemental Table 4:** Genome assembly metrics for all isolates used in this study. Column headers are as follows:

sampleName - identifier of each isolate

contigs – number of contigs per assembly

totalLength - number of all bases for all contigs for a isolate

N50 – estimate N50 values for each isolate

**Supplemental Table 5:** ST with allele numbers parsed by method for each sample. For el_gato output, the following alternative ST definitions are used:

- ’Novel ST’: all seven loci were identified, but their unique combination is not present in the ST database.
- ’Novel ST*’: at least one locus did not have an exact sequence in the database suggesting a novel allele.
- ’MA?’: multiple alleles were identified at one or more loci, preventing resolution of the true allele and assignment of an ST.
- ’MD-’: missing data for at least one locus (e.g., due to low read coverage or missing sequence in the assembly), so no ST could be determined.
- ’-’ (hyphen): missing data for this locus, often from low read coverage or absent sequence in the assembly, so no allele number could be assigned.
- ’NAT’: no match to any allele in the database, suggesting a novel allele.
- ’?’ multiple alleles identified at this locus, but the correct allele could not be resolved.

For the other two tools, all unidentified alleles or STs are simply marked with a hyphen (‘-’), meaning unknown.

Column headers are as follows:

sampleName – identifier for each isolate

ST = sequence type

*flaA –* allele number for the *flaA* gene

*pilE –* allele number for the *pilE* gene

*asd* – allele number for the *asd* gene

*mip –* allele number for the mip gene

*mompS –* allele number for the *mompS* gene

*proA –* allele number for the *proA* gene

*neuA/neuAh –* allele number for the *neuA/neuAh* gene

method *–* the tool used to determine the ST for that isolate

**Supplemental Table 6:** McNemar’s test results were used to evaluate the performance of different SBT tools compared to the Sanger-based approach. Group 1 and Group 2 columns represent the pairwise comparison of all tools. The output includes the raw p-value (p), the adjusted p-value (p.adj), and the significance level of the adjusted p-value (p.adj.signif). Adjusted p-values were calculated using the Benjamini-Hochberg method to control the false discovery rate. Statistical significance is indicated as follows: ns: not significant (p > 0.05), **: p ≤ 0.01 and > 0.001, ***: p ≤ 0.001 and > 0.0001, and ****: p ≤ 0.0001. a) This is for only the three tools (el_gato, legsta, and mompS) along with the Sanger comparator. b) This table includes all tools in table a, along with the output of using both raw and trimmed fastq data with el_gato and assemblies.

