## Supplemental Material for "Epidemiology of *Legionella*: Genome-bAsed Typing (el_gato) - a new bioinformatic tool for identifying sequence-based types of *Legionella pneumophila* from whole genome sequencing data"

**Subtitle:** Overcoming Typing Limitations in *L. pneumophila* with el_gato

**Authors:**

Alan J. Collins^*^,

Dev Mashruwala^*^,

Vasanta Chivukula^1^,

Natalia A. Kozak-Muiznieks^2^,

Lavanya Rishishwar^1^,

Emily T. Norris^1^,

Melisa J. Willby^2^,

Jennafer A. P. Hamlin^2^,

Will A. Overholt^1^,

**^1^** Applied Bioinformatics Laboratory (ASRT, Inc.; IHRC, Inc.), Atlanta, Georgia, USA

**^2^** Division of Bacterial Disease, Centers for Disease Control and Prevention, Atlanta, Georgia, USA

* Equally contributing first authors

**Input for el_gato**

el_gato can process both FASTQ read data and assembled genomes, though caution is advised with assemblies, as critical information such as read orientation is lost during assembly. This information can be important for determining the *mompS* locus. Using reads also allows el_gato to leverage read quality and coverage for applying quality control rules. In contrast, assemblies may contain errors introduced during the assembly process, which el_gato cannot detect, potentially leading to incorrect results. For paired-end reads, files can be in “.fastq” or “.fastq.gz” format, with standard differentiation between read pairs (e.g., R1/R2). el_gato does not accept single-end read data. Overall, usage is simple:

- assemblies: `el_gato.py -a <assembly file>`
- paired-end reads: `el_gato.py -1 <read1 file> -2 <read2 file>`

**Output from el_gato**
After a run, el_gato will print the identified ST of your sample to your terminal standard out and creates several files written to the specified output directory (default: out/). el_gato writes the ST profile as a tab-delimited without the headings. If you run el_gato with the `-e` flag, it includes the headings, which are displayed as such:
`Sample ST flaA pilE asd mip mompS proA neuA_neuAh`
The sample column contains the user-provided or inferred sample name. The ST column contains the overall sequence type of the sample, followed by each allele identity determined for the seven loci. The ST column can contain two kinds of values. If the identified ST corresponds to a profile found in the database, el_gato provides the corresponding number. If el_gato finds no matching ST profile or if el_gato is unable to make a confident call, then this will be reflected in the value displayed in the ST column. el_gato reports the corresponding allele number for each gene if an exact match is found in the database. Alternatively, el_gato may also note the following symbols:

| **Symbol** | **Meaning** |
| --- | --- |
| Novel ST | Novel Sequence Type: All 7 target genes found but combination is not present in the profile database. |
| Novel ST* | Novel Sequence Type due to novel allele: One or multiple target genes have a novel allele found. |
| MD- | Missing Data: ST is unidentifiable because of or more of the target genes that are unidentifiable. |
| MA? | Multiple Alleles: ST is ambiguous due to multiple alleles that could not be resolved. |
| NAT | Novel Allele Type: BLAST cannot find an exact allele match - most likely a new allele. |
| ‘-’ | Missing Data: Both percent and length identities are too low to return a match or N's in sequence. |
| ‘?’ | Multiple Alleles: More than one allele was found and could not be resolved. |

**Supplemental Methods Table 1:** Definition of alternative values for allele or ST determination

If symbols are present in the ST profile, the other output files produced by el_gato will provide additional information to understand what is being communicated.

*Description of the six output files generate by el_gato*
The files included in the output directory for a sample are as follows:

1. identified_alleles.fna
   The nucleotide sequence of all identified alleles is written in this file. If more than one allele is determined for the same locus, they are numbered arbitrarily. FASTA headers in this file correspond to the locus; if multiple copies are identified, they correspond to the query IDs from the BLAST output reported in the intermediate_outputs.txt. el_gato calls other programs to perform intermediate analyses. The outputs of those programs are provided in this file. In addition, essential log messages are also written in this file to help with troubleshooting issues. The following information may be contained in this file, depending on if the input is either read pairs or assembly:
   i. Reads-only - samtools coverage command output. See samtools coverage documentation for more information about headers at https://www.htslib.org/doc/samtools-coverage.html
   ii. Reads-only - Information about the orientation of *mompS* sequencing primer in reads mapping to biallelic sites.
   iii. BLAST output indicates the best match for identified alleles. See BLAST output documentation for more information about headers at https://www.ncbi.nlm.nih.gov/books/NBK279684/table/appendices.T.options_common_to_all_blast/
2. possible_mlsts.txt
   This file would contain all possible ST profiles if el_gato identified multiple possible alleles for any loci. In addition, if multiple *mompS* alleles were found, the information used to determine the primary allele is reported in two columns: "mompS_reads_support" and "mompS_reads_against."
   i. mompS_reads_support indicates the number of reads associated with each allele that contains the reverse sequencing primer in the expected orientation, which suggests that this is the primary allele.
   ii. mompS_reads_against indicates the number of reads containing the reverse sequencing primer in the wrong orientation and thus demonstrates that this is the secondary allele. These values are used to infer which allele is the primary *mompS* allele, and their values can be considered to represent the confidence of this characterization.
3. reads_vs_all_ref_filt_sorted.bam
   el_gato maps the provided reads to a set of reference sequences in the el_gato database (db) directory. The mapped reads are then used to extract the sequences present in the sample to identify the alleles and, ultimately, the ST. reads_vs_all_ref_filt_sorted.bam and its associated file reads_vs_all_ref_filt_sorted.bai contains the mapping information used by el_gato. The BAM file can be viewed using software such as IGV (https://software.broadinstitute.org/software/igv/) to understand better the data used by el_gato to make allele calls. Additionally, this file is a good starting point for investigating the cause of incorrectly resolved loci.
4. reads_vs_all_ref_filt_sorted.bam.bai
   Index file allows a program such as IGV to read *.bam to view the read data associated with the specific *.bam fiile.
5. report.json
   For each sample, el_gato outputs a JSON file that contains relevant information about the run, which can be included in the report PDF via the reporting script (elgato_report.py). The report script generates a pdf file with the following information dependent on if read or assembly data is included as described below. Additionally, header and footer information can be specified by the user. Please see SupplementalFigure1-elgatoBatchReport-Reads.pdf as an example of the report generated when using reads and SupplementalFigure2-elgatoBatchReport-Assemblies.pdf as an example of the report when using assemblies. For both reports, they include two pages which summarize the output (Report Summary, i) and describe relevant information (Definitions Overview, ii) as described below. After those pages, the information displayed differs dependent on the input: Paired-End Reads Report (iii) or in the Assembly Results (iv).
   i. Report Summary: Complete ST profile for every sample JSON file listed after using the `elgato_report.py` function.
   ii. Definitions Overview: Includes both ST definitions key and how the evidence for determining support of the correct *mompS* allele is assessed.
   iii. Paired-End Reads Report run specific data: ST and allele numbers parsed by an isolate’s FASTQ read data.
        i. Paired-end reads Locus Information: Information regarding each locus determined from analyzing the read data.
           1. Locus – Gene identifier
           2. Percent Covered – Percent of the locus to which reads were mapped
           3. Mean Depth – average number of reads mapped to each position within the locus.
           4. Minimum Depth – Lowest value of depth for a locus. This is based on a single base pair position within the locus.
           5. Low Depth Bases – Number of bases that fall below the depth threshold which can be specified by the user (default:10).
   iv. Assembly Results: ST and allele numbers determined based on an isolate's genome assembly.
       i. Locus Information: Information regarding each allele found in the assembly.
          1. Locus – Gene identifier
          2. Allele – Allele number for each gene. If there are multiple alleles as a possible option for a locus, that information will be listed as multiple allele numbers for that locus
          3. Contig – The contig where the gene was identified
          4. Start – The start position of the gene within the contig
          5. Stop – The stop position of the gene within the contig
          6. % Length – Percent of the length of the allele that aligned to the specified region of the contig
6. run.log
   A detailed log of the steps taken during el_gato's running includes the outputs of any programs called by el_gato and any errors encountered. Some command outputs include headers (e.g., samtools coverage and BLAST).

*el_gato report outputs for read- and assembly-based analyses*

Example outputs from the elgato_report.py function demonstrate reporting for both read-based and assembly-based analyses (Supplemental Figures 1 and 2). The read-based report includes a summary of sequence type (ST) calls and allelic profiles, a definitions overview describing abbreviations and the criteria used to determine the primary *mompS* identity, and isolate-specific pages with locus-level metrics and detailed *mompS* determination (Supplemental Figure 1). The assembly-based report presents analogous summary, definitions, and isolate-level results for the same isolates, organized under Assembly Results (Supplemental Figure 2).

**Methodology of el_gato when using genome assemblies**

Six of the seven loci (*flaA*, *pilE*, *asd*, *mip*,*proA*, and *neuA/neuAh*) are identified using BLASTn with a length threshold ('--length', default 0.3) and a percent sequence identity threshold ('--sequence', default 95.0). The length threshold specifies the minimum proportion of the query sequence that must align to the target, while the percent identity threshold defines the minimum sequence identity required to retain a BLAST hit. If multiple copies meeting these thresholds are detected, el_gato issues a warning and does not report an allele for the affected locus, which may result in no ST being assigned. Otherwise, BLASTn results are reported if they match an allele in the database. For loci that do not meet the thresholds, a symbol indicates why el_gato could not make a definitive call (Supplemental Methods Table 1). Only *mompS* requires additional processing when analyzing an assembly.

*mompS and Assemblies*

*mompS* is sometimes present in multiple copies in *Legionella pneumophila*, though typically two copies. When typing *L. pneumophila* using Sanger sequencing, primers amplify only the correct *mompS* locus. We, therefore, use *in silico* PCR (Kuhn *et al.*, 2013) to extract the correct *mompS* sequence from the assembly. The primers used for in *silico* PCR (isPCR) are mompS-430F (TTGACCATGAGTGGGATTGG) and mompS-1116R (TGGATAAATTATCCAGCCGGACTTC) as previously determined (Gaia *et al.*, 2005; Gordon *et al.*, 2017). The *mompS* allele is identified using BLASTn. If isPCR fails to detect *mompS,* standard BLASTn is used as an alternative to attempt identification. Note that this approach does not perform as well as using FASTQ reads because some information is lost during assembly. Consequently, the assembly-based method in el_gato does not apply the additional quality filtering used in the reads-based approach.

*Assembly results compared to other tools and el_gato read data*

el_gato demonstrated much reduced accuracy when using genome assemblies (SupplementalTable6-egToolComparisonToSangerResults.csv). el_gato with assemblies correctly predicted 59.2% ST calls (n = 261 out of 441). Each locus had the following correctly identified alleles as compared to the Sanger results: *flaA* with 98.6% (n = 435/441), *pilE* with 98.4% (n = 434/441), *asd* with 98.6% (n = 435/441), *mip* with 98.9% (n = 436/441), *mompS* with 59.9 % (n = 264/441), *proA* with 99.1% (n = 437/441), and *neuA/neuAh* with 98.9% (n = 436/441; Supplemental Table 5). In addition, el_gato with assemblies incorrectly typed one isolate (D7309) for the *mompS* locus. The much-reduced accuracy for determining ST based on genome assemblies is a result of an inability to correctly type the *mompS* gene. Additionally, el_gato with assemblies has a fast runtime, with an average of 1.44 seconds per assembly (minimum of 0.60 – maximum of 2.7 seconds) using the same compute resources.

**Supplemental Figure 1:** Example of the output from the elgato_report.py function when using read data. The ‘Report Summary’ page includes the ST call and allelic profile for each isolate tested. The ‘Definitions Overview’ page defines any abbreviations and how el_gato evaluates evidence for determining the primary *mompS* identity. Then the remaining pages are parsed by isolate (‘Paired-end reads report’) and include additional locus specific metrics along with a table detailing the information used to determine the *mompS* gene*.* The same isolates used to generate the reads report were also used to generate the assembly-based report.

**Supplemental Figure:** Example of the output from the elgato_report.py function. The ‘Report Summary’ page includes the ST call and allelic profile for each isolate tested. The ‘Definitions Overview’ page defines any abbreviations and how el_gato evaluates evidence for determining the primary *mompS* identity. The remaining pages are parsed by isolate (‘Assembly Results’) and include additional locus specific metrics, including whether multiple alleles were identified*.* The same isolates used to generate the reads report were also used to generate the assembly-based report.
