## SupplementalFigure1-elgatoBatchReport-Reads for "Epidemiology of *Legionella*: Genome-bAsed Typing (el_gato) - a new bioinformatic tool for identifying sequence-based types of *Legionella pneumophila* from whole genome sequencing data"

National Center for Immunization and Respiratory Diseases (NCIRD)
Division of Bacterial Diseases (DBD)/Pneumonia & Streptococcus Laboratory Branch (PSLB)
Pneumonia Response and Surveillance Laboratory (PRSL)

### **Epidemiology of *Legionella*: Genome-bAsed Typing (el_gato) Batch Results Report**

```
 /\_/\
( o.o )
 > ^ <
```

#### **Report Summary**

Sequence Based Typing (SBT) is based on 7 *Legionella pneumophila* loci (*flaA*, *pilE*, *asd*, *mip*, *mompS*, *proA*, *neuA/neuAh*). Each locus is assigned an allele number based on comparison of its sequence with sequences in an allele database. The allelic profile is the combination of allele numbers for all seven loci in order and denotes a unique Sequence Type (ST). el_gato utilizes either a genome assembly (.fasta) or Illumina paired-end reads (.fastq) to accomplish *Legionella pneumophila* SBT.More information about each sample can be found in the log file generated by el_gato. While a key to the definitions is found on the **Definition Overview** page.

| Sample ID | ST | flaA | pilE | asd | mip | mompS | proA | neuA |
|---|---|---|---|---|---|---|---|---|
| D4283_---ST_170821_Leg104-MKI-D4283-N710S508_S16_L001 | 213 | 2 | 19 | 5 | 10 | 18 | 1 | 2 |
| D4314_---ST_161115_Leg73-MKI-D4314-N703S502_S2_L001 | 1 | 1 | 4 | 3 | 1 | 1 | 1 | 1 |
| D7110_1.fastq | 59 | 7 | 6 | 17 | 3 | 13 | 11 | 11 |
| D7309_---ST_170728_Leg101-MKI-D7309-N711S503_S3_L001 | 213 | 2 | 19 | 5 | 10 | 18 | 1 | 2 |

This test has not been cleared or approved by the FDA. The performance characteristics have been established by the Pneumonia and Streptococcus Laboratory Branch. The results are intended for public health purposes only and must NOT be communicated to the patient, their care provider, or placed in the patient's medical record. These results should NOT be used for diagnosis, treatment, or assessment of patient health or management.
Reference Value: Not applicable.

#### Definitions Overview

##### ST Definitions Key

Novel ST = the alleles for all 7 loci were identified, however their unique combination and corresponding ST has not been found in the database.

Novel ST* = an exact match for sequences of at least one locus was not identified in the database, which may indicate a novel allele.

MA? = **m**ultiple **a**lleles; for at least one locus, multiple alleles were identified, and the true allele could not be resolved; therefore, no ST could be determined.

MD- = **m**issing **d**ata; data were missing for at least one locus (e.g., low read coverage at one or more position, missing sequence in assembly); therefore, no ST could be determined.

'-' = missing data; data were missing for this locus (e.g., low read coverage at one or more position, missing sequence in assembly); therefore, an allele number could not be determined.

'NAT' = **n**ovel **a**llele **t**ype; this locus did not match any allele listed in the database, possibly indicating a novel allele.

'?' = multiple alleles; for this locus multiple alleles were identified, and could not be resolved.

##### Evidence for support of *mompS* allele call

Multiple *mompS* alleles may be identified in the genome of a single *L. pneumophila* isolate. In these instances, it is important to use the correct mompS allele to generate the allelic profile and ST. el_gato resolves this issue by considering the orientation of a small "primer" sequence in relation to the reference sequence. Please see the el_gato README for more details.

"NA" indicates that primer support was not assessed since only one mompS allele was identified. Otherwise, the primary mompS allele is identified using the following criteria:

1. Only one allele has associated reads with the correctly oriented primer.

2. One allele has more than 3 times as many reads with the correctly oriented primer as the other.

3. One allele has no associated reads with the primer in either orientation, but the other has reads with the primer only in the wrong direction. The sequence with no associated reads is considered the primary locus in this case.

4. Absence of any primer-associated reads does not allow identification of the primary allele.

This test has not been cleared or approved by the FDA. The performance characteristics have been established by the Pneumonia and Streptococcus Laboratory Branch. The results are intended for public health purposes only and must NOT be communicated to the patient, their care provider, or placed in the patient's medical record. These results should NOT be used for diagnosis, treatment, or assessment of patient health or management.
Reference Value: Not applicable.

**Epidemiology of *Legionella*: Genome-bAsed Typing (el_gato) Paired-End Reads Report**

#### D4283_---ST_170821_Leg104-MKI-D4283-N710S508_S16_L001 reads report

The following sample was analyzed using the paired-end reads functionality with el_gato version 1.20.1. The tables below show the full ST profile of the sample, the coverage data for each locus, and information regarding the primers used to identify the primary mompS allele. Low depth bases indicate bases that do not have 10 or more reads covering that base, unless the default depth cutoff was adjusted. More information can be found in the log file for this sample.

| Sample ID | ST | flaA | pilE | asd | mip | mompS | proA | neuA |
|---|---|---|---|---|---|---|---|---|
| D4283_---ST_170821_Leg104-MKI-D4283-N710S508_S16_L001 | 213 | 2 | 19 | 5 | 10 | 18 | 1 | 2 |

##### Locus Information

| Locus | Percent Covered | Mean Depth | Minimum Depth | Low depth bases |
|---|---|---|---|---|
| flaA | 100.0 | 158.5 | 128.0 | 0.0 |
| pilE | 100.0 | 90.1 | 68.0 | 0.0 |
| asd | 100.0 | 101.9 | 60.0 | 0.0 |
| mip | 100.0 | 77.5 | 56.0 | 0.0 |
| mompS | 100.0 | 263.9 | 203.0 | 0.0 |
| proA | 100.0 | 98.6 | 59.0 | 0.0 |
| neuA | 100.0 | 74.5 | 36.0 | 0.0 |

##### *mompS* Primer Information

| Allele | Reads Indicating Primary | Reads Indicating Secondary |
|---|---|---|
| mompS_63 | 0 | 0 |
| mompS_18 | 28 | 0 |

Please find the key for definitions and evidence for support of *mompS* allele call on the Definitions Overview page.

This test has not been cleared or approved by the FDA. The performance characteristics have been established by the Pneumonia and Streptococcus Laboratory Branch. The results are intended for public health purposes only and must NOT be communicated to the patient, their care provider, or placed in the patient's medical record. These results should NOT be used for diagnosis, treatment, or assessment of patient health or management.
Reference Value: Not applicable.

#### D4314_---ST_161115_Leg73-MKI-D4314-N703S502_S2_L001 reads report

The following sample was analyzed using the paired-end reads functionality with el_gato version 1.20.1. The tables below show the full ST profile of the sample, the coverage data for each locus, and information regarding the primers used to identify the primary mompS allele. Low depth bases indicate bases that do not have 10 or more reads covering that base, unless the default depth cutoff was adjusted. More information can be found in the log file for this sample.

| Sample ID | ST | flaA | pilE | asd | mip | mompS | proA | neuA |
|---|---|---|---|---|---|---|---|---|
| D4314_---ST_161115_Leg73-MKI-D4314-N703S502_S2_L001 | 1 | 1 | 4 | 3 | 1 | 1 | 1 | 1 |

##### Locus Information

| Locus | Percent Covered | Mean Depth | Minimum Depth | Low depth bases |
|---|---|---|---|---|
| flaA | 100.0 | 181.9 | 151.0 | 0.0 |
| pilE | 100.0 | 130.3 | 89.0 | 0.0 |
| asd | 100.0 | 138.5 | 116.0 | 0.0 |
| mip | 100.0 | 126.8 | 105.0 | 0.0 |
| mompS | 100.0 | 341.6 | 280.0 | 0.0 |
| proA | 100.0 | 147.8 | 71.0 | 0.0 |
| neuA | 100.0 | 79.7 | 61.0 | 0.0 |

##### *mompS* Primer Information

| Allele | Reads Indicating Primary | Reads Indicating Secondary |
|---|---|---|
| mompS_1 | NA | NA |

Please find the key for definitions and evidence for support of *mompS* allele call on the Definitions Overview page.

This test has not been cleared or approved by the FDA. The performance characteristics have been established by the Pneumonia and Streptococcus Laboratory Branch. The results are intended for public health purposes only and must NOT be communicated to the patient, their care provider, or placed in the patient's medical record. These results should NOT be used for diagnosis, treatment, or assessment of patient health or management.
Reference Value: Not applicable.

#### D7110_1.fastq reads report

The following sample was analyzed using the paired-end reads functionality with el_gato version 1.20.1. The tables below show the full ST profile of the sample, the coverage data for each locus, and information regarding the primers used to identify the primary mompS allele. Low depth bases indicate bases that do not have 10 or more reads covering that base, unless the default depth cutoff was adjusted. More information can be found in the log file for this sample.

| Sample ID | ST | flaA | pilE | asd | mip | mompS | proA | neuA |
|---|---|---|---|---|---|---|---|---|
| D7110_1.fastq | 59 | 7 | 6 | 17 | 3 | 13 | 11 | 11 |

##### Locus Information

| Locus | Percent Covered | Mean Depth | Minimum Depth | Low depth bases |
|---|---|---|---|---|
| flaA | 100.0 | 140.2 | 112.0 | 0.0 |
| pilE | 100.0 | 84.7 | 61.0 | 0.0 |
| asd | 100.0 | 91.8 | 67.0 | 0.0 |
| mip | 100.0 | 70.9 | 58.0 | 0.0 |
| mompS | 100.0 | 245.2 | 207.0 | 0.0 |
| proA | 100.0 | 94.4 | 58.0 | 0.0 |
| neuA | 100.0 | 70.2 | 53.0 | 0.0 |

##### *mompS* Primer Information

| Allele | Reads Indicating Primary | Reads Indicating Secondary |
|---|---|---|
| mompS_13 | NA | NA |

Please find the key for definitions and evidence for support of *mompS* allele call on the Definitions Overview page.

This test has not been cleared or approved by the FDA. The performance characteristics have been established by the Pneumonia and Streptococcus Laboratory Branch. The results are intended for public health purposes only and must NOT be communicated to the patient, their care provider, or placed in the patient's medical record. These results should NOT be used for diagnosis, treatment, or assessment of patient health or management.
Reference Value: Not applicable.

**Epidemiology of *Legionella*: Genome-bAsed Typing (el_gato) Paired-End Reads Report**

#### D7309_---ST_170728_Leg101-MKI-D7309-N711S503_S3_L001 reads report

The following sample was analyzed using the paired-end reads functionality with el_gato version 1.20.1. The tables below show the full ST profile of the sample, the coverage data for each locus, and information regarding the primers used to identify the primary mompS allele. Low depth bases indicate bases that do not have 10 or more reads covering that base, unless the default depth cutoff was adjusted. More information can be found in the log file for this sample.

| Sample ID | ST | flaA | pilE | asd | mip | mompS | proA | neuA |
|---|---|---|---|---|---|---|---|---|
| D7309_---ST_170728_Leg101-MKI-D7309-N711S503_S3_L001 | 213 | 2 | 19 | 5 | 10 | 18 | 1 | 2 |

##### Locus Information

| Locus | Percent Covered | Mean Depth | Minimum Depth | Low depth bases |
|---|---|---|---|---|
| flaA | 100.0 | 153.9 | 124.0 | 0.0 |
| pilE | 100.0 | 139.1 | 101.0 | 0.0 |
| asd | 100.0 | 151.3 | 124.0 | 0.0 |
| mip | 100.0 | 121.1 | 92.0 | 0.0 |
| mompS | 100.0 | 373.8 | 283.0 | 0.0 |
| proA | 100.0 | 147.4 | 63.0 | 0.0 |
| neuA | 100.0 | 118.0 | 67.0 | 0.0 |

##### *mompS* Primer Information

| Allele | Reads Indicating Primary | Reads Indicating Secondary |
|---|---|---|
| mompS_63 | 0 | 0 |
| mompS_18 | 37 | 0 |

Please find the key for definitions and evidence for support of *mompS* allele call on the Definitions Overview page.

This test has not been cleared or approved by the FDA. The performance characteristics have been established by the Pneumonia and Streptococcus Laboratory Branch. The results are intended for public health purposes only and must NOT be communicated to the patient, their care provider, or placed in the patient's medical record. These results should NOT be used for diagnosis, treatment, or assessment of patient health or management.
Reference Value: Not applicable.
