## SupplementalFigure2-elgatoBatchReport-Assemblies for "Epidemiology of *Legionella*: Genome-bAsed Typing (el_gato) - a new bioinformatic tool for identifying sequence-based types of *Legionella pneumophila* from whole genome sequencing data"

**Epidemiology of *Legionella*: Genome-bAsed Typing (el_gato) Batch Results Report**

```
 /\_/\
( o.o )
 > ^ <
```

### Report Summary

Sequence Based Typing (SBT) is based on 7 *Legionella pneumophila* loci (*flaA*, *pilE*, *asd*, *mip*, *mompS*, *proA*, *neuA/neuAh*). Each locus is assigned an allele number based on comparison of its sequence with sequences in an allele database. The allelic profile is the combination of allele numbers for all seven loci in order and denotes a unique Sequence Type (ST). el_gato utilizes either a genome assembly (.fasta) or Illumina paired-end reads (.fastq) to accomplish *Legionella pneumophila* SBT.More information about each sample can be found in the log file generated by el_gato. While a key to the definitions is found on the **Definition Overview** page.

| Sample ID | ST | flaA | pilE | asd | mip | mompS | proA | neuA |
|---|---|---|---|---|---|---|---|---|
| D4283_---ST_170821_L... | MA? | 2 | 19 | 5 | 10 | ? | 1 | 2 |
| D4314_---ST_161115_L... | MD- | 1 | 4 | 3 | 1 | - | 1 | 1 |
| D7110 | 59 | 7 | 6 | 17 | 3 | 13 | 11 | 11 |
| D7309_---ST_170728_L... | Novel ST | 2 | 19 | 5 | 10 | 63 | 1 | 2 |

This test has not been cleared or approved by the FDA. The performance characteristics have been established by the Pneumonia and Streptococcus Laboratory Branch. The results are intended for public health purposes only and must NOT be communicated to the patient, their care provider, or placed in the patient's medical record. These results should NOT be used for diagnosis, treatment, or assessment of patient health or management.
Reference Value: Not applicable.

### D4283 ---ST 170821 Leg104-MKI-D4283-N710S508 S16 genomic report

The following sample was analyzed using the assembly functionality with el_gato version 1.21.0. The tables below show the full ST profile of the sample and the corresponding locus location information. Unless specified by the user, el_gato utilizes a default 30% (0.3) BLAST hit length threshold and a 95% (95.0) sequence identity threshold to identify the presence of multiple copies of an allele. el_gato will only report allele matches for BLAST hits of 100% length and 100% identity. More information can be found in the log file for this sample.

| Sample ID | ST | flaA | pilE | asd | mip | mompS | proA | neuA |
|---|---|---|---|---|---|---|---|---|
| D4283_---ST_170821_L... | MA? | 2 | 19 | 5 | 10 | ? | 1 | 2 |

### Locus Information

| locus | allele | contig | start | stop | %length |
|---|---|---|---|---|---|
| flaA | flaA_2 | NODE_1_length_402422_cov_... | 279320 | 279501 | 100.0 |
| pilE | pilE_19 | NODE_17_length_75691_cov_... | 23333 | 23665 | 100.0 |
| asd | asd_5 | NODE_12_length_130689_cov... | 122689 | 123161 | 100.0 |
| mip | mip_10 | NODE_2_length_340674_cov_... | 180924 | 181325 | 100.0 |
| mompS | mompS_63 | NODE_7_length_162959_cov_... | 69521 | 69872 | 100.0 |
| | mompS_18 | NODE_7_length_162959_cov_... | 70556 | 70754 | 56.5 |
| proA | proA_1 | NODE_10_length_135509_cov... | 125390 | 125794 | 100.0 |
| neuA_neuAH | neuA_neuAH_2 | NODE_2_length_340674_cov_... | 137303 | 137656 | 100.0 |

% Length = BLAST hit length as a percent of expected locus size.

This test has not been cleared or approved by the FDA. The performance characteristics have been established by the Pneumonia and Streptococcus Laboratory Branch. The results are intended for public health purposes only and must NOT be communicated to the patient, their care provider, or placed in the patient's medical record. These results should NOT be used for diagnosis, treatment, or assessment of patient health or management.
Reference Value: Not applicable.

### D4314 ---ST 161115 Leg73-MKI-D4314-N703S502 S2 genomic report

The following sample was analyzed using the assembly functionality with el_gato version 1.21.0. The tables below show the full ST profile of the sample and the corresponding locus location information. Unless specified by the user, el_gato utilizes a default 30% (0.3) BLAST hit length threshold and a 95% (95.0) sequence identity threshold to identify the presence of multiple copies of an allele. el_gato will only report allele matches for BLAST hits of 100% length and 100% identity. More information can be found in the log file for this sample.

| Sample ID | ST | flaA | pilE | asd | mip | mompS | proA | neuA |
|---|---|---|---|---|---|---|---|---|
| D4314_---ST_161115_L... | MD- | 1 | 4 | 3 | 1 | - | 1 | 1 |

#### Locus Information

| locus | allele | contig | start | stop | %length |
|---|---|---|---|---|---|
| flaA | flaA_1 | NODE_2_length_399372_cov_... | 225256 | 225437 | 100.0 |
| pilE | pilE_4 | NODE_18_length_75769_cov_... | 23412 | 23744 | 100.0 |
| asd | asd_3 | NODE_19_length_71844_cov_... | 7515 | 7987 | 100.0 |
| mip | mip_1 | NODE_17_length_79249_cov_... | 31108 | 31509 | 100.0 |
| mompS | mompS_99 | NODE_5_length_239040_cov_... | 94778 | 94906 | 36.6 |
| proA | proA_1 | NODE_6_length_220013_cov_... | 93989 | 94393 | 100.0 |
| neuA_neuAH | neuA_neuAH_1 | NODE_17_length_79249_cov_... | 72886 | 73239 | 100.0 |

% Length = BLAST hit length as a percent of expected locus size.

This test has not been cleared or approved by the FDA. The performance characteristics have been established by the Pneumonia and Streptococcus Laboratory Branch. The results are intended for public health purposes only and must NOT be communicated to the patient, their care provider, or placed in the patient's medical record. These results should NOT be used for diagnosis, treatment, or assessment of patient health or management.
Reference Value: Not applicable.

### D7110 genomic report

The following sample was analyzed using the assembly functionality with el_gato version 1.21.0. The tables below show the full ST profile of the sample and the corresponding locus location information. Unless specified by the user, el_gato utilizes a default 30% (0.3) BLAST hit length threshold and a 95% (95.0) sequence identity threshold to identify the presence of multiple copies of an allele. el_gato will only report allele matches for BLAST hits of 100% length and 100% identity. More information can be found in the log file for this sample.

| Sample ID | ST | flaA | pilE | asd | mip | mompS | proA | neuA |
|---|---|---|---|---|---|---|---|---|
| D7110 | 59 | 7 | 6 | 17 | 3 | 13 | 11 | 11 |

### Locus Information

| locus | allele | contig | start | stop | %length |
|---|---|---|---|---|---|
| flaA | flaA_7 | NODE_3_length_209247_cov_... | 48524 | 48705 | 100.0 |
| pilE | pilE_6 | NODE_30_length_34642_cov_... | 9693 | 10025 | 100.0 |
| asd | asd_17 | NODE_6_length_153525_cov_... | 91338 | 91810 | 100.0 |
| mip | mip_3 | NODE_14_length_85849_cov_... | 22935 | 23336 | 100.0 |
| proA | proA_11 | NODE_4_length_193370_cov_... | 47440 | 47844 | 100.0 |
| neuA_neuAH | neuA_neuAH_11 | NODE_22_length_61886_cov_... | 18796 | 19149 | 100.0 |

% Length = BLAST hit length as a percent of expected locus size.

This test has not been cleared or approved by the FDA. The performance characteristics have been established by the Pneumonia and Streptococcus Laboratory Branch. The results are intended for public health purposes only and must NOT be communicated to the patient, their care provider, or placed in the patient's medical record. These results should NOT be used for diagnosis, treatment, or assessment of patient health or management.
Reference Value: Not applicable.

### D7309 ---ST 170728 Leg101-MKI-D7309-N711S503 S3 genomic report

The following sample was analyzed using the assembly functionality with el_gato version 1.21.0. The tables below show the full ST profile of the sample and the corresponding locus location information. Unless specified by the user, el_gato utilizes a default 30% (0.3) BLAST hit length threshold and a 95% (95.0) sequence identity threshold to identify the presence of multiple copies of an allele. el_gato will only report allele matches for BLAST hits of 100% length and 100% identity. More information can be found in the log file for this sample.

| Sample ID | ST | flaA | pilE | asd | mip | mompS | proA | neuA |
|---|---|---|---|---|---|---|---|---|
| D7309_---ST_170728_L... | Novel ST | 2 | 19 | 5 | 10 | 63 | 1 | 2 |

#### Locus Information

| locus | allele | contig | start | stop | %length |
|---|---|---|---|---|---|
| flaA | flaA_2 | NODE_1_length_402422_cov_... | 279320 | 279501 | 100.0 |
| pilE | pilE_19 | NODE_17_length_75051_cov_... | 52027 | 52359 | 100.0 |
| asd | asd_5 | NODE_18_length_72052_cov_... | 7529 | 8001 | 100.0 |
| mip | mip_10 | NODE_2_length_341470_cov_... | 159350 | 159751 | 100.0 |
| proA | proA_1 | NODE_10_length_135509_cov... | 9716 | 10120 | 100.0 |
| neuA_neuAH | neuA_neuAH_2 | NODE_2_length_341470_cov_... | 203026 | 203379 | 100.0 |

% Length = BLAST hit length as a percent of expected locus size.

This test has not been cleared or approved by the FDA. The performance characteristics have been established by the Pneumonia and Streptococcus Laboratory Branch. The results are intended for public health purposes only and must NOT be communicated to the patient, their care provider, or placed in the patient's medical record. These results should NOT be used for diagnosis, treatment, or assessment of patient health or management.
Reference Value: Not applicable.
