## SupplementalTable2-SBT_primers for "Epidemiology of *Legionella*: Genome-bAsed Typing (el_gato) - a new bioinformatic tool for identifying sequence-based types of *Legionella pneumophila* from whole genome sequencing data"

**Supplemental Table 2**

**SBT Primers**

| **Loci** | **Primer Name** | **Sequence 5'-3'** | **amplicon size** | **Primer type** |
| --- | --- | --- | --- | --- |
| flaA | flaA-587F | **TGTAAAACGACGGCCAGT** GCG TAT TGC TCA AAA TAC TG | 428 | amplification |
|  | flaA-960R | **CAGGAAACAGCTATGACC** CCA TTA ATC GTT AAG TTG TAG G |  |  |
| pilE | pilE-35F | **TGTAAAACGACGGCCAGT** CAC AAT CGG ATG GAA CAC AAA CTA | 477 | amplification |
|  | pilE-453R | **CAGGAAACAGCTATGACC** GCT GGC GCA CTC GGT ATC T |  |  |
| asd | asd-511F | **TGTAAAACGACGGCCAGT** CCC TAA TTG CTC TAC CAT TCA GAT G | 588 | amplification |
|  | asd-1039R | **CAGGAAACAGCTATGACC** CGA ATG TTA TCT GCG ACT ATC CAC |  |  |
| mip | mip-74F | **TGTAAAACGACGGCCAGT** GCT GCA ACC GAT GCC AC | 573 | amplification |
|  | mip-595R | **CAGGAAACAGCTATGACC** CAT ATG CAA GAC CTG AGG GAA C |  |  |
| mompS | mompS-450F | **TGTAAAACGACGGCCAGT** TTG ACC ATG AGT GGG ATT GG | 732 | amplification |
|  | mompS-1126R | TGG ATA AAT TAT CCA GCC GGA CTT C |  |  |
| proA | proA-1107F | **TGTAAAACGACGGCCAGT** GAT CGC CAA TGC AAT TAG | 499 | amplification |
|  | proA-1553R | **CAGGAAACAGCTATGACC** ACC ATA ACA TCA AAA GCC |  |  |
| neuA | neuA-196F | **TGTAAAACGACGGCCAGT** CCG TTC AAT ATG GGG CTT CAG | 471 | amplification |
|  | neuA-611R | **CAGGAAACAGCTATGACC** CGA TGT CGA TGG ATT CAC TAA TAC |  |  |
| flaA alternate^1^ | flaA-LN | TAT GCG TGA GCT TTC CGT TC | 493 | Alternative amplification primers |
|  | flaA-RN | GGT ATC ACC TGC GGT TCC A |  |  |
| neuAh^1^ | neuA-left2 | ATC CAG CAG TTT TTA MAA ATT TAG G | 791-794 | Alternative amplification primers |
|  | neuA-right2 | TGG CTG CAT AAA YTA ATT CTT TAG CCA |  |  |
| M13 tag | M13F | TGT AAA ACG ACG GCC AGT | n/a | Sequencing primer |
| M13 tag | M13R | CAG GAA ACA GCT ATG ACC | n/a | Sequencing primer |
| mompS | MompS-1015R | CAG AAG CTG CGA AAT CAG | n/a | Sequencing primer |

**^1^** When alternative amplification flaA and neuA primers are used (flaA-LN, flaA-RN, neuA-left2, and neuA-right2), the alternative amplification primers are also used for sequencing.
