## SupplementalTable6McNemarOutput for "Epidemiology of *Legionella*: Genome-bAsed Typing (el_gato) - a new bioinformatic tool for identifying sequence-based types of *Legionella pneumophila* from whole genome sequencing data"

**Supplemental Table 6**

McNemar test results comparing ST assignment agreement between tools (including alternative el_gato inputs) and Sanger sequencing.

| **Comparison** | **p-value** | **Adjusted p-value** | **Significance** |
| --- | --- | --- | --- |
| eg1.21.0Assembly vs eg1.21.1Raw | 2.70 × 10⁻³⁹ | 3.78 × 10⁻³⁸ | **** |
| eg1.21.0Assembly vs eg1.21.1Trimmed | 2.70 × 10⁻³⁹ | 3.78 × 10⁻³⁸ | **** |
| eg1.21.0Assembly vs legsta | 2.66 × 10⁻¹⁵ | 3.72 × 10⁻¹⁴ | **** |
| eg1.21.0Assembly vs mompS | 2.15 × 10⁻³³ | 3.01 × 10⁻³² | **** |
| eg1.21.0Assembly vs Sanger | 2.18 × 10⁻⁴⁰ | 3.05 × 10⁻³⁹ | **** |
| eg1.21.1Raw vs eg1.21.1Trimmed | NaN | NaN | – |
| eg1.21.1Raw vs legsta | 7.07 × 10⁻⁵⁶ | 9.90 × 10⁻⁵⁵ | **** |
| eg1.21.1Raw vs mompS | 1.77 × 10⁻⁴ | 2.48 × 10⁻³ | ** |
| eg1.21.1Raw vs Sanger | 7.36 × 10⁻² | 1.00 × 10⁰ | ns |
| eg1.21.1Trimmed vs legsta | 7.07 × 10⁻⁵⁶ | 9.90 × 10⁻⁵⁵ | **** |
| eg1.21.1Trimmed vs mompS | 1.77 × 10⁻⁴ | 2.48 × 10⁻³ | ** |
| eg1.21.1Trimmed vs Sanger | 7.36 × 10⁻² | 1.00 × 10⁰ | ns |
| legsta vs mompS | 2.18 × 10⁻⁵² | 3.05 × 10⁻⁵¹ | **** |
| legsta vs Sanger | 5.75 × 10⁻⁵⁷ | 8.05 × 10⁻⁵⁶ | **** |
| mompS vs Sanger | 1.27 × 10⁻⁵ | 1.78 × 10⁻⁴ | *** |

ns = not signifcant; ** = p < 0.01; *** = p < 0.001; **** = p < 0.0001; NaN = identical results
